# Microbiological, biochemical, physicochemical surface properties and biofilm forming ability of *Brettanomyces bruxellensis*

**DOI:** 10.1101/579144

**Authors:** Maria Dimopoulou, Margareth Renault, Marguerite Dols-Lafargue, Warren Albertin-Leguay, Jean-Marie Herry, Marie-Noëlle Bellon-Fontaine, Isabelle Masneuf-Pomarede

**Affiliations:** UR Oenologie EA 4577, USC 1366 INRA, Bordeaux INP, Université de Bordeaux, France; UMR Génie et Microbiologie des Procédés Alimentaires, GMPA, AgroParisTech, INRA, Université Paris-Saclay, 78850 Thiverval-Grignon, France; ENSCBP, Bordeaux INP, 33600 Pessac, France; Ecole Nationale Supérieure des Sciences Agronomiques de Bordeaux-Aquitaine, France

**Author notes:** Corresponding author: Maria Dimopoulou.

**Keywords:** *Brettanomyces bruxellensis*, wine spoilage, biochemical properties, physico-chemical surface properties, biofilm

## Abstract

*Brettanomyces bruxellensis* is a serious source of concern for winemakers. The production of volatile phenols by the yeast species confers to wine unpleasant sensory characteristics which are unacceptable by the consumers and inevitably provoke economic loss for the wine industry. This ubiquitous yeast is able to adapt to all winemaking steps and to withstand various environmental conditions. Moreover, the ability of *B. bruxellensis* to adhere and colonize inert materials can be the cause of the yeast persistence in the cellars and thus recurrent wine spoilage. We therefore investigated the surface properties, biofilm formation capacity and the factors which may affect the attachment of the yeast cells to surfaces with eight strains representative of the genetic diversity of the species. Our results show that the biofilm formation ability is strain-dependent and suggest a possible link between the physicochemical properties of the studied strains and their corresponding genetic group.

## Introduction

The process of fermentation has been used for years to improve the shelf-life and the sensory properties of food and beverages. Even if nowadays, the fermentation is often carried out from monocultures of a single fermentative strain, the traditional process usually involves a multitude of different microorganisms naturally present in the environment (Liu et al. 2017). These microorganisms produce a vast variety of aromatic molecules, and while some of them ameliorate the sensory profile and satisfy the consumers, some others cause sensory defects and product rejection (Belda et al. 2017; Tempère et al. 2018). The *B. bruxellensis* species may belong to both categories, depending on the production process and the final product characteristics. This yeast species can be desirable in kombucha and beer production, while in wine, it is considered as the source of major organoleptic defects (Chatonnet et al. 1992, 1995; Steensels et al. 2015). Actually, the capacity of *B. bruxellensis* to produce volatile phenols, that impart a “stable” or “horse-sweat” odor, can cause a drop in wine quality and in some cases the product becomes unfit for sale (Agnolucci et al. 2017).

Recently, the genome sequencing of *B. bruxellensis* demonstrated a great intraspecific genetic diversity and in particular, a variability of the ploidy level (Fournier et al. 2017; Borneman et al. 2014). A rapid, reliable and discriminating method has been developed as a tool for genetic typing of *B. bruxellensis* strains using specific microsatellite markers (Albertin et al. 2014; Avramova et al. 2018a). Based on this method, a great number of *B. bruxellensis* isolates from various niches (beer, wine, cider, bioethanol, kombucha) and geographical areas were genotyped. The population structure analysis revealed three main genetic clusters and three sub-clusters associated with the strains ploidy level and substrates of isolation. Interestingly, the strain tolerance/resistance to sulfur dioxide, the main antimicrobial compound used in wine differs from one genetic cluster to the other, unraveling a strong link between genotype and this phenotypic trait (Avramova et al. 2018a, b). Other methods have been developed to control the spoilage yeast growth, as for example inactivation by heat or pressure, sterilizing filtration or through the use of ionized radiation. However these techniques are often expensive and may have a negative impact on sensory properties, which makes them incompatible with the production of high quality wines (Gerland 2010). In addition, the majority of the current techniques do not systematically eliminate the entire population of *B. bruxellensis*. This situation may be explained by the high intraspecific genetic and phenotypic diversity observed within the yeast species that differentiates some strains and increases their adaptation mechanism. Indeed, the repeated use of high doses of the antimicrobial agent sulphur dioxyde to control the development of *B. bruxellensis* during winemaking, may have led to the emergence of more resistant strains which can tolerate the current used doses (Avramova et al. 2018b; Capozzi et al. 2016; Curtin et al. 2012; Conterno et al. 2010; Dimopoulou et al. 2019).

This yeast is able to survive and multiply in wine, especially in case of sluggish fermentation and difficulties of other species to monopolize the wine ecosystem (Renouf et al. 2006; Romano et al. 2008). Even if *B. bruxellensis* was detected at low level on grape berry, repetitive infections were observed in wine, suggesting that the cellar, rather than the vineyard, could be the main source of contamination (Barata et al. 2008; Garde-Cerdan and Ancin-Azpilicueta 2006; Gonzales-Arenzana et al. 2013; Rubio et al. 2015). Despite the fact that wine is not produced continuously throughout the year, yeast persistence in the cellars was demonstrated from year to year (Grangeteau et al. 2016). However, the mechanism by which yeasts persist in the winery is not yet elucidated.

Yeast cells possess a remarkable ability to adhere to abiotic surfaces, in particular in response to stress and nutrient limitation (Verstrepen and Klis 2006). Adhesion to surfaces and subsequent biofilm formation enable long-term survival of fungi and bacteria in unfavorable nutrient environments (Tek et al. 2018). Moreover, disinfectants are not able to penetrate the biofilm matrix and possibly the resistance to anti-microbial compounds or cleaning agents could be related to the ability of microorganism to form biofilm (Carpentier and Cerf 1993; Perpetuini et al. 2018). Yeast adhesion is one of the most plastic and variable examined phenotype, while the attachment ability of closely genetic related strain could vary dramatically (Verstrepen and Klis 2006). Adhesion to abiotic surfaces is the first step in biofilm formation and depends on the physico-chemical properties of cells as well as of materials surfaces. The cell surface properties are linked to the molecular composition of the wall and other external elements of the micro-organisms. In *Saccharomyces cerevisiae*, the attachment to plastic and mat formation requires Flo11p, a member of the large family of fungal cell surface glycoproteins (Reynolds and Fink 2001). The same authors showed that mat formation and cell architecture structure is modified as yeast ploidy level increases. Surface proteins such as adhesins can also increase the cell-surface hydrophobicity and promote hydrophobic interaction between the cells and abiotic surfaces (Kang and Choi 2005).

Joseph et al. (2007) showed that the majority of the *B. bruxellensis* isolates studied were able to produce biofilm onto polystyrene surface, upon long incubation time in the presence of low sugar concentration. However, this first study did not consider the genetic diversity of the species, nor the growth rate and yield of the studied strains.

In this paper, various methods were tested to examine the microbiological, biochemical, physicochemical surface properties and biofilm forming ability of a panel of strains of *B. bruxellensis*, representative of the species genetic diversity. A putative correlation between these different properties and the strain genotype has been examined.

## Materials and Methods

### Strains and growth conditions

*B. bruxellensis* isolates were obtained from a variety of regions and fermented substrates, being part of the CRB Oenologie collection (Centre de Ressources Biologiques Œnologie, Bordeaux, France). The eight strains used in this study and their genetic group (Avramova et al. 2018a) are listed in Table 1. Their distribution on the genetic dendrogram is presented in Fig. 1. Firstly, all strains were spotted onto YPD medium containing 1% (w/v) yeast extract (Difco Laboratories, Detroit,), 2% (w/v) Bacto peptone (Difco), and 1% (w/v) glucose and agar (20g/L) and incubated for 5 days at 25°C. One cm^2^ of solid culture was then inoculated into 40 mL sterile Erlenmeyer flasks containing liquid YPD broth, placed at 25°C at 150 rpm, for 48h and then transferred into 250 mL liquid YPD culture for 10 days until the stationary growth phase, collected at density from 2,89 (strain L0417) to 4 (strain CBS 2499 and AWRI1608).

**Table 1.**
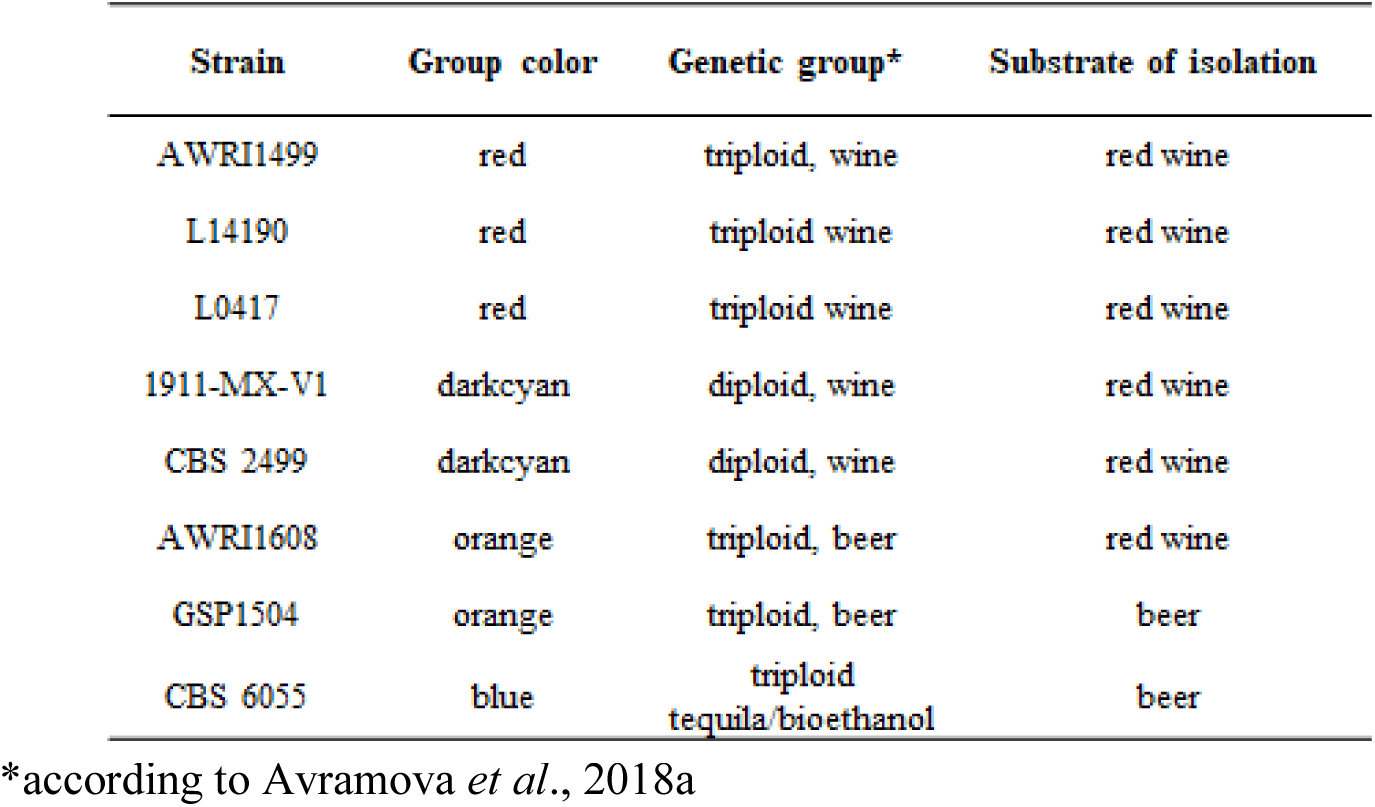
The eight *B. bruxellensis* strains used in this study

| Strain | Group color | Genetic group* | Substrate of isolation |
| --- | --- | --- | --- |
| AWRI1499 | red | triploid, wine | red wine |
| L14190 | red | triploid wine | red wine |
| L0417 | red | triploid wine | red wine |
| 1911-MX-V1 | darkcyan | diploid, wine | red wine |
| CBS 2499 | darkcyan | diploid, wine | red wine |
| AWRI1608 | orange | triploid, beer | red wine |
| GSP1504 | orange | triploid, beer | beer |
| CBS 6055 | blue | triploid<br>tequila/bioethanol | beer |

\*according to Avramova *et al.*, 2018a

**Fig. 1.**
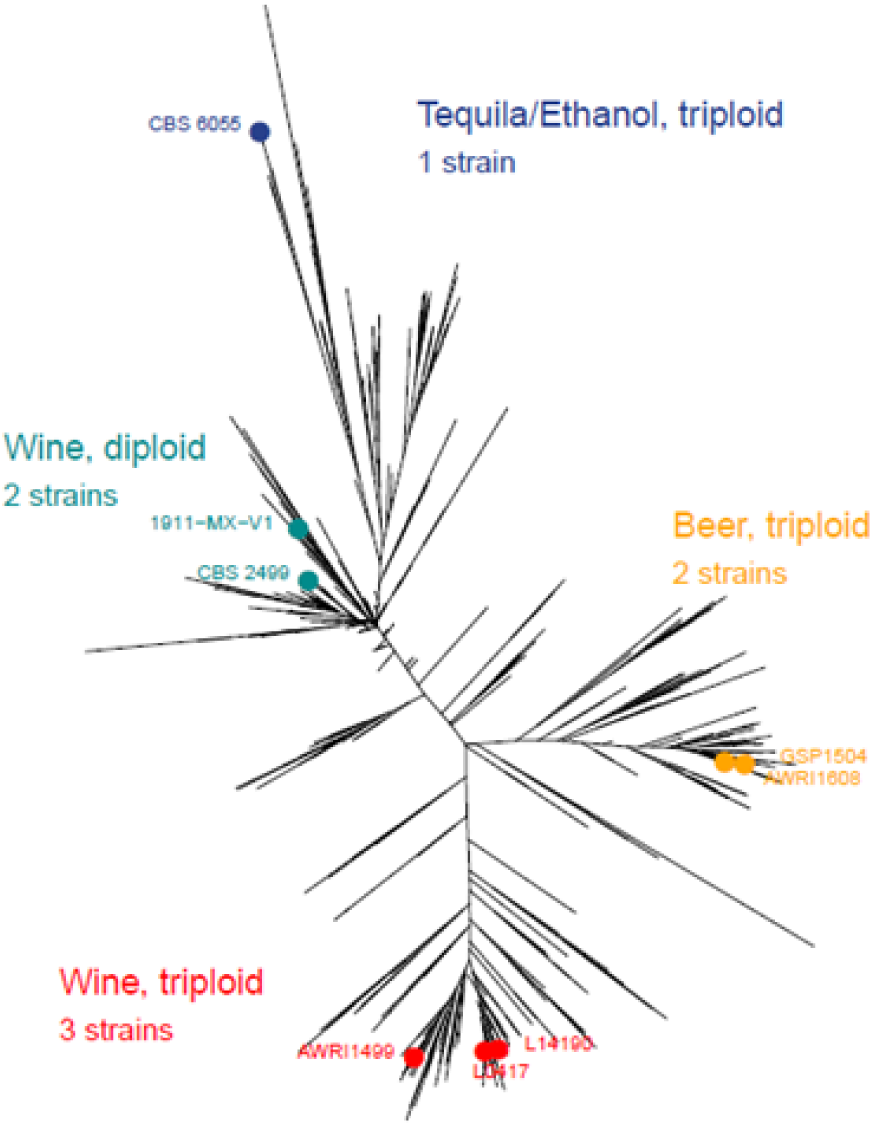
Position of the eight *B. bruxellensis* strains in a dendrogram of 1488 strains (Avramova et al. 2018a)

### Microbiological characteristics

#### Mat formation

Yeast strains were evaluated for their ability for mat formation as described by Reynolds and Fink (2001), Briefly, 10 µL of liquid culture in YPD were spotted onto YPD with 1.5% (w/v) agar. The plates were then incubated for 17 days at 25°C and photographed using wide angle digital camera (Nikon, Coolpix P500). Analyses were performed in duplicate except for the strain GSP1504.

#### Biofilm biomass quantification to polystyrene microtiter plates

Biofilm formation was quantified using a colorimetric microtiter 96 well polystyrene plate (Thermo Scientific Nunc MicroWell) by a method adapted from O’Toole et al. (2000) (Tek et al. 2018). The eight strains were grown in YPD medium for 10 days until stationary growth phase and then inoculated in YPD or in wine like medium (45% Sauvignon juice, 50% water and 5% ethanol).The OD of inoculum was spectrophotometrically adjusted to 0.1 at 600 nm in order to calibrate all the strains at a similar initial population level and cells were suspended in YPD or in wine like medium. The inoculated media were aliquoted (200 μL) in polystyrene microplate wells. Each strain was inoculated in eight replicate wells in two separate plates, while eight negative control wells containing only broth were also included in each plate. The plates were incubated without any agitation at 25°C for 15 days in a hermetic plastic box containing a glass of water to create a highly humid atmosphere and avoid evaporation. After 15 days of incubation, the suspended cells population was estimated by optical absorbance (OD 600 nm) by using a single well in which the broth was mixed with a micropipette. The other wells were gently washed twice with 200 µL of 0.9 % (v/v) NaCl, dried in an inverted position and stained with 1% (w/v) crystal violet. The wells were rinsed again and the crystal violet was solubilized in 100µL of ethanol:acetone (80:20, v/v). The absorbance at 595 nm was determined using a microplate reader (Molecular devices).

### Biochemical characteristics

#### Cell lipid extraction

After 10 days of growth in YPD medium, cells were collected and a sample corresponding to 2x10^9^ cells was withdrawn. Cell concentration was estimated by flow cytometry and optical density (OD 600nm). Then each sample was washed twice with distilled water, frozen with liquid nitrogen and conserved at -20°C in Eppendorf tubes. From the frozen samples obtained previously, the membrane lipid extraction was realized according to the protocol of Tronchoni et al. (2012) with some slight modifications. In each tube, 1 g of glass beads (0.5 mm, Biospec Products), 700 μL of cold methanol and 140 μL of EDTA 0.1M were added and vigorously mixed (4 x 45 s) in a mini-bead-beater-8 (Biospec Products, Qiagen). By this way, the cell membrane wall was completely disordered and the liquid phase was separated from the glass beads by centrifugation and transferred into a 15 mL glass screw tube. Lipid extraction was performed in three steps, starting with the addition of 2.5 mL chloroform/methanol (1:1, v/v, for 45 min), centrifugation (3000 rpm, 5 min) and recuperation of the inferior phase. Then 2.5 mL of chloroform/methanol (2:1, v/v, for 45 min) were added and the inferior phase was recuperated as before. The last extraction was realized by adding 2.5 mL chloroform/methanol (1:2, v/v, for 45 min). The inferior organic phases was transferred to a 15 mL glass screw tube and cleaned twice by adding KCl 0.88% (1/4 of a total volume of the extract). After vortexing and cooling at 4°C for 30 min, the samples were centrifuged (3000 rpm, 5 min) and the inferior organic phase was collected and stored at -80°C until analysis.

#### Fatty acid determination

The extracted lipids were concentrated to dryness under nitrogen stream and then methylated by the presence of methanol and sulphuric acid (5% v/v) for 2 h at 85°C in the presence of 50 μg heptanoic acid (Sigma Aldrich) as internal standard. Then the addition in the same tube of 1 mL of NaCl (2.5% w/v) and 2 mL of hexane enabled the extraction of methylic esters from the fatty acids, which were concentrated to dryness under nitrogen stream. Finally 200 μL of hexane were added to fatty acids extract and their relative concentrations were determined by Gas Chromatography coupled with Flame Ionisation, as described by Redon et al. (2009).

#### Exopolysaccharides quantification

After 10 days of growth in YPD broth, soluble exopolysaccharides (EPS) liberated by the yeasts were collected in the culture supernatant. The total excreted polysaccharides were precipitated (Dimopoulou et al. 2016) and their concentration was determined according to the anthrone-sulfuric method with glucose as standard (Ludwig and Goldberg 1956). For each sample, the polymer precipitation and assays were done in triplicate. The EPS quantification results were normalized by the cell population after 10 days of growth (OD 600 nm).

### Surface physicochemical properties

#### Preparation of yeasts suspensions characteristics

The physicochemical characterization of *B. bruxellensis* strains was carried out for cells grown in YPD broth. Briefly, cells were harvested by centrifugation (Eppendorf) at 4°C for 10 min at 7000 g and then washed twice with and re-suspended in the relevant suspending liquid (NaCl 150mM or 1.5 mM). All experiments were performed on three separately grown cultures.

#### Measurement of electrophoretic mobility (EM)

The electrophoretic mobility (EM) of yeast in a sodium chloride solution (1.5 mM) was measured as a function of the pH within the range of 2 to 5, adjusted by the addition of HNO_3_. The concentration of the suspension was approximately at 10^7^ cells/mL. Measurements were taken in a 50 V/cm electric field with a laser zetameter (CAD Instruments, France). For each measurement, results were based on the automated video analysis of about 200 cells. Each experiment was performed twice.

#### Microbial adhesion to hydrocarbon

Microbial adhesion to hydrocarbon (MATH) enables the evaluation of the hydrophobic/hydrophilic character of the cell surface of *B. bruxellensis* strains (Bellon-Fontaine et al. 1996). Experimentally, yeast suspension (1.5 mL) was mixed with 0.25 mL of each solvent (hexadecane, decane). The mixture was stirred for 2 min to form an emulsion and a rest period of 15 min allowed the complete separation of the two phases. The optical density (OD) of the aqueous phase and that of the initial cell suspension (OD_0_) were measured at 400 nm. The microbial affinity to each solvent was calculated using the formula:

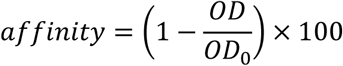

Each experiment was performed twice.

### Statistical analysis

Kruskal-Wallis statistical test (agricolae package, R, p-value < 0.05) and Principal Component Analysis (PCA) were performed using R-package.

## Results

### Microbiological characteristics

Eight strains of *B. bruxellensis* belonging to distinct genetic groups were used: two diploid strains (belonging to the genetic group dark cyan, CBS2499_like), three triploid strains (red group AWRI1499_like), two triploid strains (orange group AWRI1608_like) and a triploid tequila / bioethanol group strain (blue group, CBS5512 like) (Table 1, Fig. 1).

The mat formation of the eight strains was observed after 17 days on YPD plate (Fig. 2). The diameter of the mats varied between 9 mm (1911-MX-V1) and 15 mm (CBS 6055). All the other strains displayed mats with a diameter ranged from 11 to 13 mm. CBS 6055 was the only strain exhibiting mat with rough edges. No relation between ploidy level and preliminary mat formation was observed The *B. bruxellensis* strains were then studied for their biofilm forming ability in polystyrene microtiter plates, in two different media, YPD and a Sauvignon must derived wine-like medium. As shown in Fig. 3, the medium composition changed significantly the biofilm formation ability of the strains belonging to the darkcyan, red and blue genetic groups. More precisely all the strains in these groups, expressed increased biofilm capacity in wine-like medium compared with in YPD broth. On the contrary, the two strains in the orange genetic group showed a similar performance in wine-like and YPD broth. Furthermore, in wine-like medium, the strain CBS 2499 displayed the highest biofilm formation ability whereas CBS 6055 displayed the lowest biofilm formation ability.

**Fig. 2.**
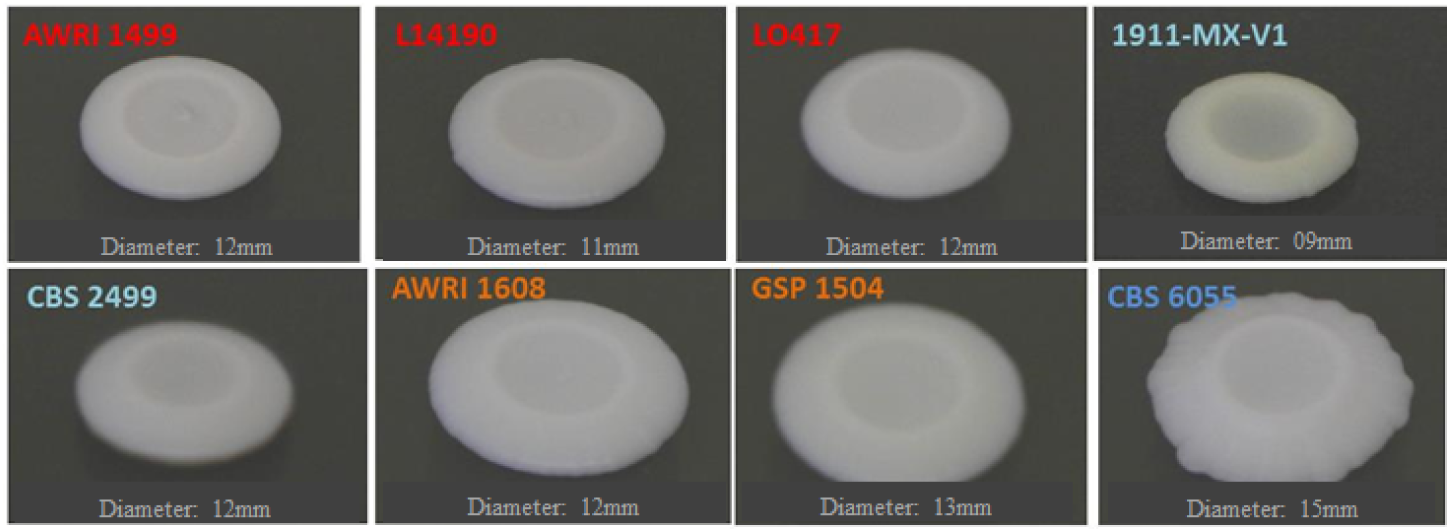
Photos of colony morphology taken by camera of the eight tested strains of *B. bruxellensis* after 17 days of growth at YPD plate.

**Fig. 3.**
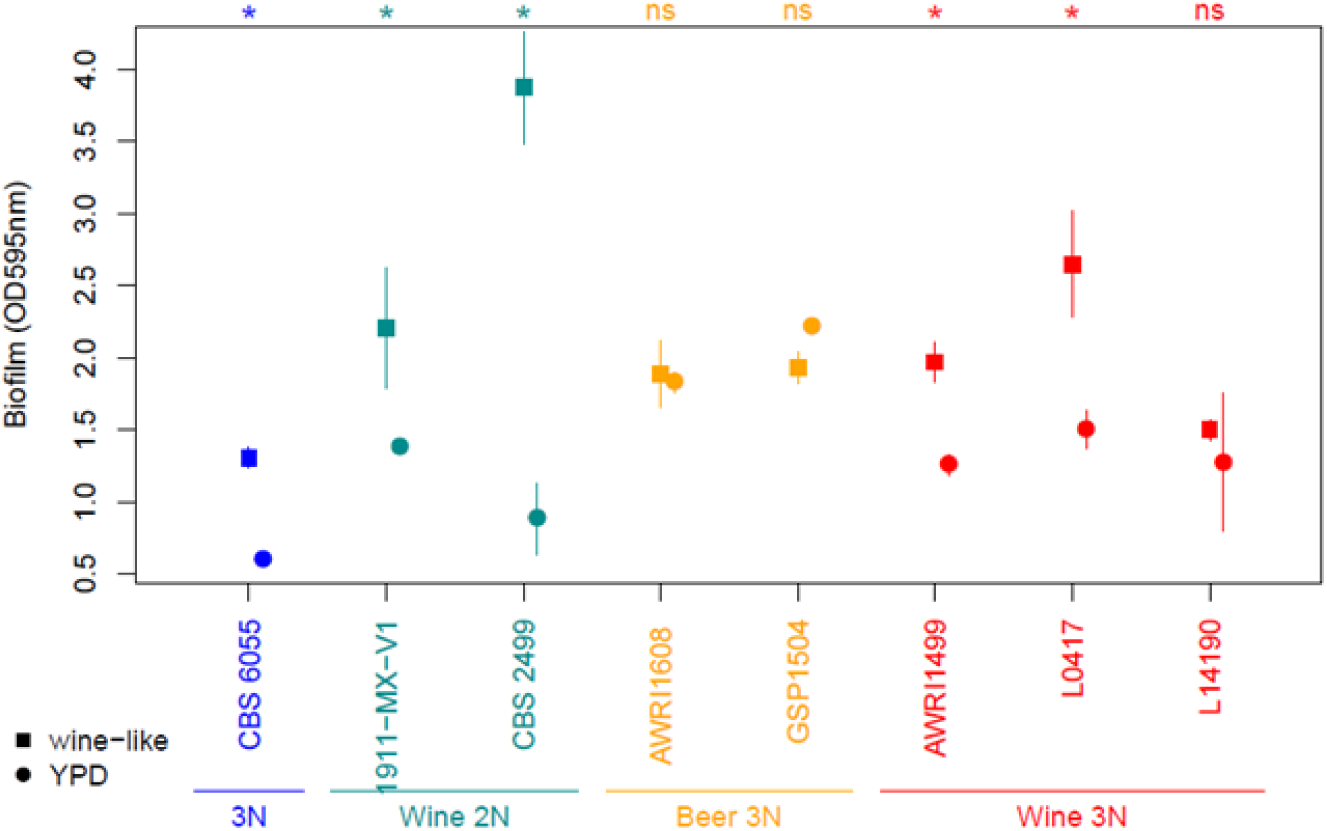
Biofilm formation ability of the eight *B. bruxellensis* strains in two different growth media; wine like and YPD. The colors represent the genetic group of the strains. Upper stars or “NS” denote respectively significant difference between media or “non-significance” as defined by Kruskal-Wallis statistical test (agricolae package, R, p-value < 0.05).

### Biochemical characteristics

#### Fatty acids composition

Fatty acids content was also studied in the eight selected strains. The total fatty acids content was in the same order of magnitude for the eight strains. However the strains differed by the fatty acid proportions. Interestingly, the SFA (Saturated Fatty Acids) to MUFA (Monounsaturated Fatty Acids) ratio was similar for each genetic subpopulation studied (Fig. 4). The highest content of SFAs, 91% of the total membrane fatty acids amount, was observed for the strains L14190, L0417 and AWRI1499. These three strains belonging to the red genetic group were significantly different from most of the strains in the other groups, with a mean SFA/MUFA ratio more than twice higher than the ratio of the other three genetic groups. On the contrary, the strain AWRI1608 showed the lowest ratio level, composed at 73% of SFA.

**Fig. 4.**
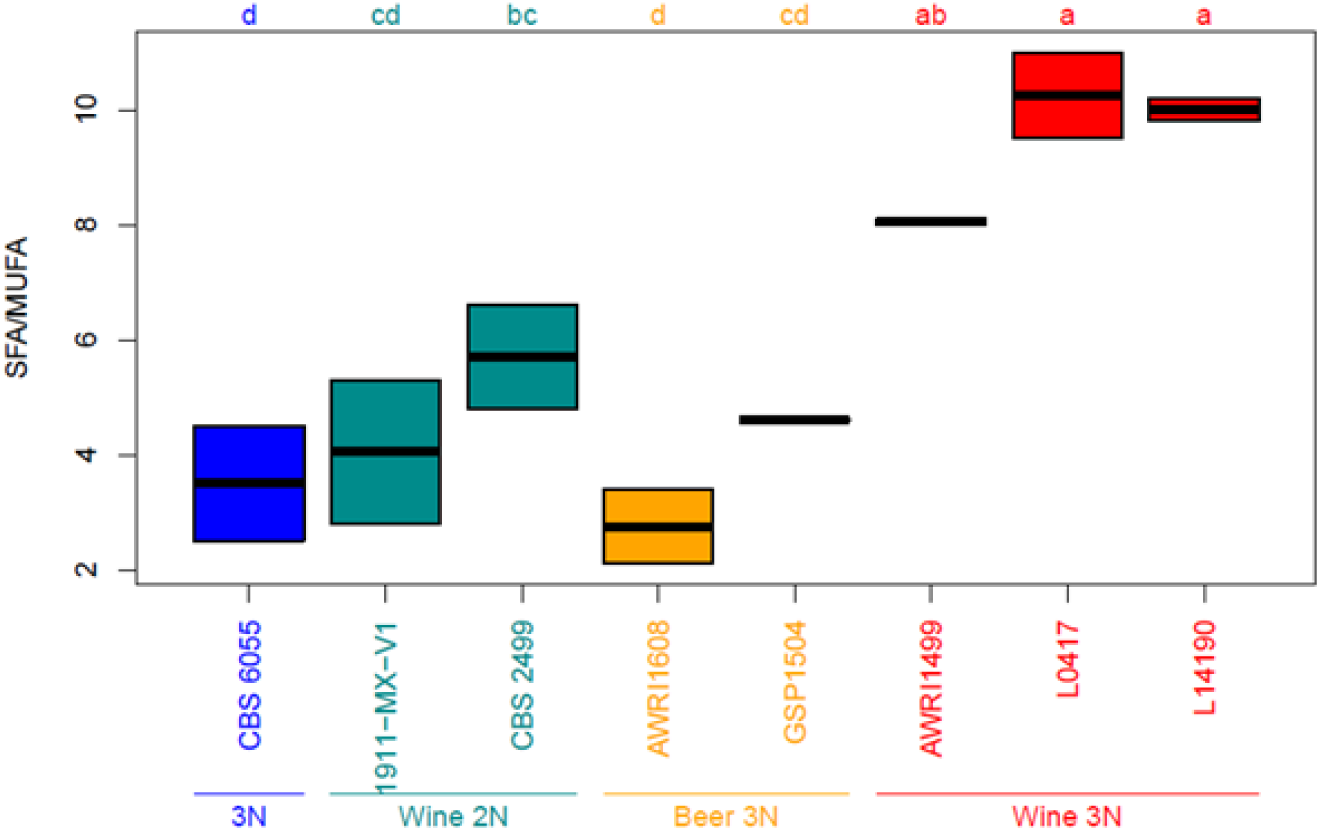
Ratio of Saturated Fatty Acid to Monounsaturated Fatty Acid of the eight tested strains of *B. bruxellensis*. The colors represent the genetic group of the strains. Upper letters represent significance groups as defined by Kruskal-Wallis statistical test (agricolae package, R, p-value < 0.05).

#### Exopolysaccharides production

The *B. bruxellensis* strains were examined for their ability to liberate exopolysaccharides (EPS) after growth in YPD medium for 10 days until reaching stationary growth phase. The results are presented as the ratio of total amount of EPS (mg/L) produced to the cell population (OD at 600 nm) (Fig. 5). The best-producing strains were CBS 6055 and L0417, which liberated more than 100 mg/L/OD of soluble exopolysaccharides in the culture medium. On the other hand, the strain 1911-MX-V1 liberated less than 40 mg/L/OD of EPS. Even if the EPS production ability of the eight studied strains could not be significantly distinguished according to their genetic group, the strains of the darkcyan genetic group displayed the lower EPS liberating ability, with a mean production of 51.5 mg/L/OD.

**Fig. 5.**
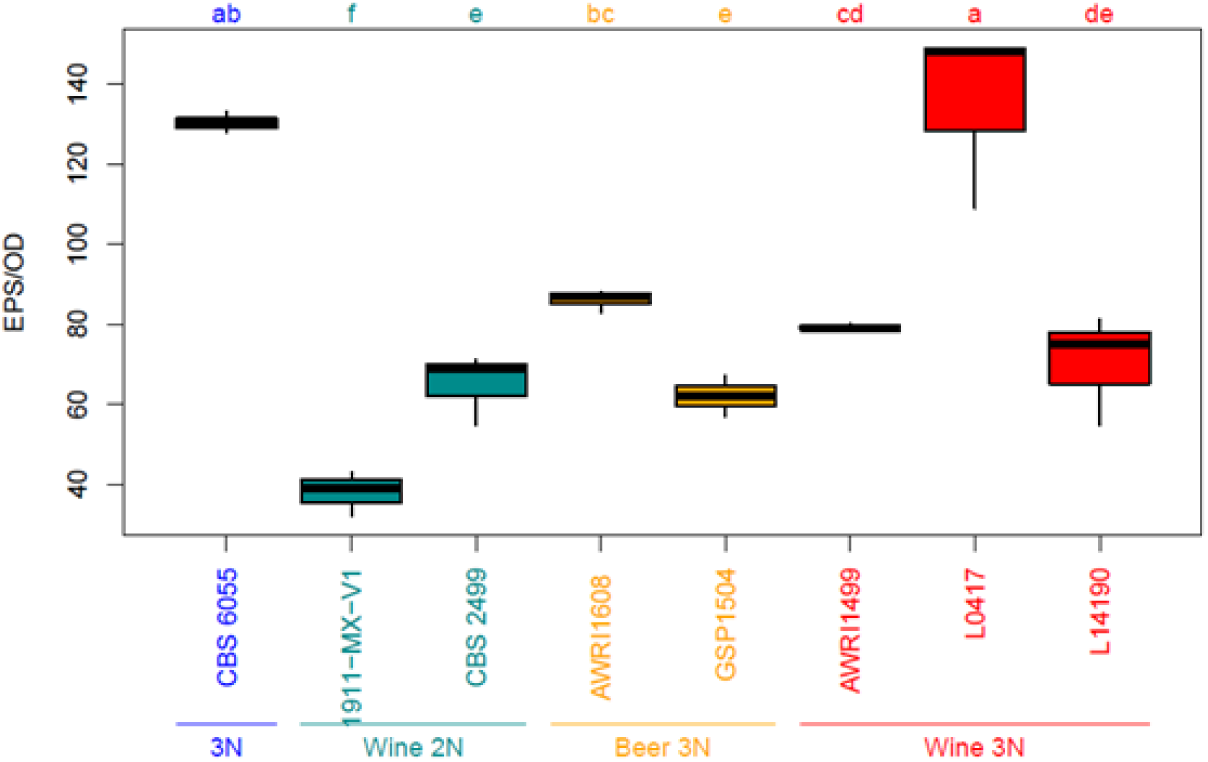
Exopolysaccharide production of the eight tested strains of *B. bruxellensis*. The colors represent the genetic group of the strains. Upper letters represent significance groups as defined by Kruskal-Wallis statistical test (agricolae package, R, p-value < 0.05).

### Surface physicochemical characteristics

The electrophoretic mobilities (EM) of the eight *B. bruxellensis* strains were measured at seven different pH values (from 2 to 5). The EM values of each strain suspended in 1.5 mM NaCl revealed negatively charged cells at pH values between 2 and 5 (Fig. 6). No isoelectric point could be determined within the range of pH values investigated but a reduction in mobility could be observed at pH 2. In our experimental conditions and whatever the tested pH, AWRI 1608, GSP 1504 and CBS 6055 (orange and blue genetic group) exhibited greater electronegativity than AWRI 1499, L0417, L14190, CBS 2499 and 1911-MX-V (red and darkcyan genetic group). Moreover, from an oenological point of view, in wine conditions (pH close to 3.5), the two strains from the orange genetic group were the less electronegative.

**Fig. 6.**
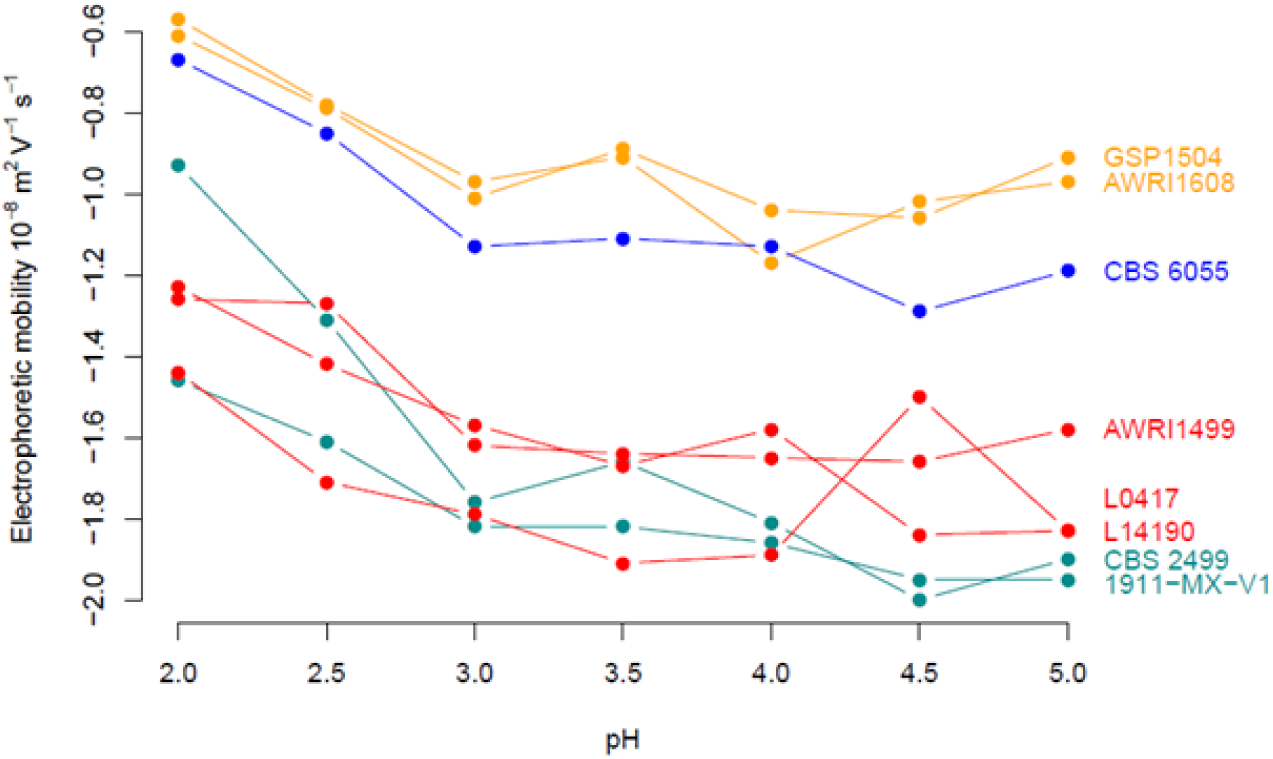
Electrophoretic mobility according to the pH of the eight strains of *B. bruxellensis*. The colors represent the genetic group of the strains. The standard deviations of EM do not exceed 0.2 for all yeasts.

The eight *B. bruxellensis* strains were also assayed for their affinity to apolar solvents, decane and hexadecane, with the MATH analysis test. According to our results the affinity of *B. bruxellensis* strains for hexadecane or decane, values ranged from 0 to 25.6% (Table 2) reflecting hydrophilic surfaces characteristics. It could be noted that in the darkcyan group, only CBS 2499 strain displayed the higher affinity for both solvents.

**Table 2.**
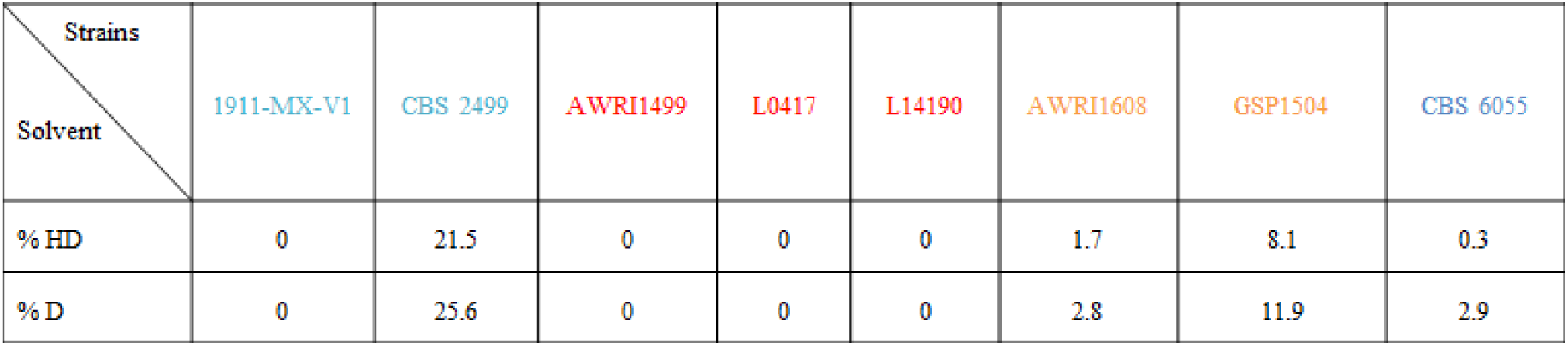
Percentage of affinity to the hexadecane and decane used in the MATH analysis for the eight *B. bruxellensis* strains studied. The standard deviations of % affinity do not exceed 0.5.

| Strains |  |  |  |  |  |  |  |  |
| --- | --- | --- | --- | --- | --- | --- | --- | --- |
| Solvent | 1911-MX-V1 | CBS 2499 | AWRI1499 | L0417 | L14190 | AWRI1608 | GSP1504 | CBS 6055 |
| % HD | 0 | 21.5 | 0 | 0 | 0 | 1.7 | 8.1 | 0.3 |
| % D | 0 | 25.6 | 0 | 0 | 0 | 2.8 | 11.9 | 2.9 |

### PCA

Principal-component analysis of combined data is shown in Fig.7. In this representation, the abscissa represented 38.5% of the total variation from the original data set and was mainly correlated with a lower negative charge of yeast cells and the mat formation. The ordinate, which represented 29.6% of the total variation from the original data set was mainly correlated with EPS production. This axis was also negatively correlated with yeast cell affinity for nonpolar solvents as well as biofilm formation in wine. This Principal-component analysis clearly distinguishes the yeast strains according to their genetic group.

**Fig. 7.**
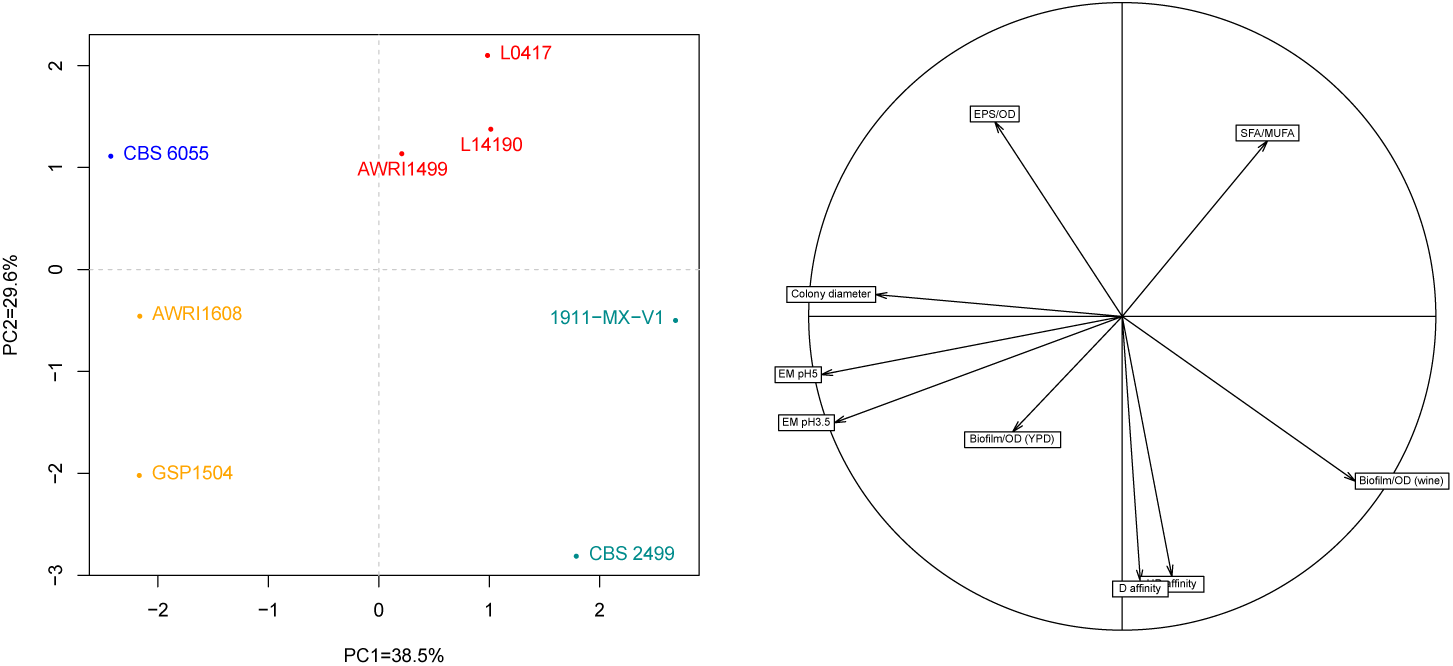
Principal-component analysis of combined data.

## Discussion

In this study, different protocols were developed and applied in order to examine the microbiological, biochemical, physicochemical surface properties and biofilm forming ability of a panel of strains of *B. bruxellensis*, representative of the species genetic diversity. Considering the high intra-species genetic diversity of *B. bruxellensis*, our study considered isolates representative of the different genetic groups of the species (Avramova et al. 2018a).

Different mat diameters and aspect were reported for the first time. By using isogenic strains from haploid to tetraploid of *Saccharomyces*, Reynolds and Fink (2001) reported an inverse relation between ploidy level and mat formation. No such relationship has been observed in this study. However mat diameter has been correlated with the yeast surface charge related to the genetic group. In the same way and depending on the genetic group, the biofilm formation into polystyrene wells was differently affected by the growth medium. *Saccharomyces cerevisiae* biofilm forming ability was shown to be linked to glucose levels, with a reduction in complete absence of glucose (Reynolds and Fink 2001). This was shown to depend to *FLO11* transcription and glucose repression (Gagiano et al. 1999). In our experimental condition, glucose but also ethanol levels were higher in wine like medium compared to YPD, and two strains (1911-MX-V1 and CBS 2499) showed higher biofilm forming ability in the medium with the highest glucose concentration. However, opposite results were obtained for the two strains of the orange genetic group as biofilm formation ability was not significantly modulated by medium composition as noticed for the other strains, thus suggesting different adhesion or regulation mechanisms according to the genetic group considered.

Interestingly, this is the first time that the membrane total free fatty acids composition and the exopolysaccharide liberation capacity of the species have been studied. The total free fatty acids membrane composition has been used in the past as a typing method (Rozes et al. 1992) and was shown to pay an important role in the cell permeability and adaptation mechanism especially under hostile environmental conditions like wine. Genes involved in lipid metabolism were showed to be enriched in *B. bruxellensis* genome with some genes that may contribute to ethanol tolerance of the species (Woolfit et al. 2007). Recent study showed that the presence of sulfites leads to increased cell permeability (Longin et al. 2016). The three strains of the red genetic group are composed of a higher ratio of SFA/MUFA compared to the other genetic groups. Taking into consideration that this genetic group gathered mainly tolerant/resistant strains to high concentration of sulphur dioxide (Avramova et al. 2018a), a link between the two tested phenotypic traits but also with the genetic group could be suggested. Regarding end-point exopolysaccharide liberation, the strains CBS 6055 and L0417 displayed the highest production capacity, which may be due to either distinct mannoprotein composition or cell wall dynamics or to premature cell lysis compared to other strains. Differences are also noted concerning the physicochemical properties of the eight yeasts analyzed which all presented hydrophilic and negatively charged profiles. Indeed, EM measurements clearly indicate a greater electronegativity of the strains of the orange and blue genetic group in comparison with strains of red or darkcyan genetic group. All these results suggest distinct wall composition and metabolism traits which can possibly affect the biofilm production capacity. Indeed, the wall polysaccharides are the first and most abundant component of the cell which comes in contact with the surfaces and can affect the microbial colonization ability (Ghafoor *et al*., 2011; Sheppard and Howell 2016). According to previous studies, they may contribute positively or negatively to biofilm formation (Legras et al. 2016; Verstrepen and Klis 2006). From a general point of view, even if the strains number was low, our present study confirmed the fact that *B. bruxellensis* shows a great variability not only at genetic but also phenotypic level. The tested strains are clearly different regarding their cell surface properties and this may have significant consequences, firstly on their ability to primarily adhere to surfaces and secondly on their biofilm formation capacity. Additional work with a higher number of strains representative of the genetic groups and a close examination of the biofilm formation steps is now needed. The detection and identification of strains contaminating the cellar and the prediction of their persistence abilities seems to be an indispensable tool for the winemakers in order to better adapt their winemaking techniques and especially their cleaning procedures.

## Acknowledgments

This work was supported by funds from FranceAgriMer. Lipids analysis was performed at the lipidomic plateform of Bordeaux University.

